# Effects of carbon to nitrogen ratio (C:N) on water quality and growth performance of *Litopenaeus vannamei* (Boone, 1931) in the biofloc system with a salinity of 5‰

**DOI:** 10.1101/2021.12.28.474292

**Authors:** Hai-Hong Huang, Chao-Yun Li, Tao Liang, Yan-Ju Lei, Pin-Hong Yang

## Abstract

This study aimed to investigate the effects of carbon to nitrogen ratio (C:N) on the water quality and shrimp growth performance during the grow-out culture of *Litopenaeus vannamei* in the biofloc system under a low salinity condition. Three biofloc treatments with an C:N (contained in the inputted feed and carbon source with the assumption that 75% of the feed nitrogen is excreted) of 8:1 (CN8), 16:1 (CN16) and 24:1 (CN24), respectively, were designed to stocking shrimp juveniles (~ 0.8 g) at a density of 270 individuals m^-3^, for a 63-days culture experiment at a salinity of about 5‰. Results showed that in CN8 treatment, the levels of pH (6.9±0.1), carbonate alkalinity (104.0±2.8mg L^-1^ CaCO_3_), biofloc volume (4.8±0.9mL L^-1^) and TSS (327.4±24.4mg L^-1^) were significantly lower than those in the other two treatments (≥7.6±0.3, ≥157.6±21.6mg L^-1^ CaCO_3_, ≥24.1±3.7mL L^-1^ and ≥508.1±32.3mg L^-1^, *P*<0.05); whereas the levels of TAN (7.1±0.9mg L^-1^), nitrite (14.0±3.6mg L^-1^) and nitrate (77.0±5.0mg L^-1^) were significantly higher than those in the other treatments (≤2.0±0.6mg L^-1^, ≤4.9±3.1mg L^-1^ and ≤14.7±5.9mg L^-1^, *P*<0.05). The zootechnical parameters of shrimp were not significantly different between three treatments (*P*>0.05), except that the survival rates in CN16 treatment (96.8±2.0%) and CN24 treatment (93.7±4.2%) were significantly higher than that of CN8 treatment (81.5±6.4%, *P*<0.05). The results indicated that an inputted C:N higher than 16:1 was suitable for the biofloc system with a low salinity of 5‰, with an optimal inferred C:N range of 18.5-21.0:1 for water quality and growth performance.

## 1. Introduction

Biofloc technology is a recent developed approach for intensive aquaculture which could assimilate inorganic nitrogen in situ to keep a good water quality without a large amount of water exchange (Avnimelech, 2015). In a biofloc system, the management of inorganic nitrogen accumulation is based upon carbon metabolism and nitrogen-immobilizing microbial processes (Avnimelech, 1999). During this process, while bacteria immobilize inorganic nitrogen as food to product new cells, carbon is also required (Deng et al., 2018; Hargreaves, 2006). However, carbon is not enough in aquaculture water body in general (Avnimelech, 1999). In a word, extra matter rich in carbon (carbohydrate or carbon source) should be added in the biofloc system (Avnimelech, 1999). After addition of the exogenous carbon source, the growth of microorganisms is prompted, accompanying enhancement of *in-situ* assimilation of nitrogen compounds accumulated in aquaculture waterbody, consequently leading to maintenance of a good water quality without substantial water exchange (Avnimelech, 2015; Ebeling et al., 2006).

The addition amount of carbon source needed to reduce/assimilate inorganic nitrogen depends on the microbial conversion coefficient, carbon to nitrogen ratio (C:N) in the microbial biomass, and carbon contents of the added material (Avnimelech, 1999). For example, assuming that the C:N in the microbial biomass, the carbon content of the added carbohydrate and the microbial conversion efficiency are 4, 50% and 0.4, respectively, the carbohydrate addition needed to reduce the total ammonia nitrogen (TAN) concentration by 1 g N is 20 g (Avnimelech, 1999).

The amount of added carbon source could also be defined as the inputted C:N (contained in all inputted materials, such as feed and carbon source). For instance, assuming 30% protein is contained in feed pellets (4.65% N) and 50% of the feed nitrogen is excreted, the inputted C:N should be raised to 15.75 (Avnimelech, 1999). In fact, different carbon contents in carbon sources, as well as the different protein contents in feeds for different species or different stage of aquatic animals, lead to that a wide-range C:N (9:1-27:1) could be suitable for biofloc systems (Chakrapani et al., 2021; Dauda et al., 2018; Liu et al., 2019; Miao et al., 2017; Panigrahi et al., 2019; Panigrahi et al., 2018; Vilani et al., 2016; Xu et al., 2016; Xu et al., 2018). Additionally, different C:N demands for the growth of different microorganism groups also make the efficient operation of biofloc systems possible under the wide-range C:N condition. For instance, with increasing of C:N, different predominant microorganism communities would be formed, such as photoautotrophic microalgaes, autotrophic nitrifiers and heterotrophic bacteria, (Becerra-Dorame et al., 2014; Ebeling et al., 2006; Haveman et al., 2009; Jung et al., 2017; Ray et al., 2011b; Schrader et al., 2011; Schveitzer et al., 2017; Xu et al., 2016). Therefore, non-proper C:N in a certain biofloc system might impact water quality, biofloc biomass production and eventually the feed utilization and shrimp performance (Avnimelech, 1999; Ebeling et al., 2006), indicating that selection of a proper C:N is very important to a biofloc system under a certain condition (Avnimelech, 2015; Tong et al., 2020).

The Pacific white-leg shrimp (*Litopenaeus vannamei*, Boon, 1931), the most widely cultivated euryhaline shrimp species in the world (FAO, 2018), could be raised in near-freshwater system at a salinity as low as 0.5‰ in inland zones (Van Wyk et al., 1999). However, culture of this marine species in inland zones might negatively affect the local ecosystem, even under the low salinity condition (Avnimelech, 2015). Recently, biofloc technology has been successfully applied in culture of *L. vannamei* at a low salinity with minimal or zero water exchange (Lobato et al., 2019; Ray and Lotz, 2017; Ray et al., 2011a), indicating that the environmental effects of culturing this marine species in inland zones would be minimized by adopting this technology (Avnimelech, 2015).

It was reported that the toxicity of ammonia and nitrite to shrimp reinforced with decreasing of salinity (Lin and Chen, 2001; Lin and Chen, 2003; Ray and Lotz, 2017). Furthermore, salinity vibrating towards low levels might change the dynamics and the presence of natural bacteria, and the predominance of specific microorganism groups adapted to the environment established in marine salinity condition (Luis et al., 2018; Morais et al., 2020; Reis et al., 2019), leading to different effectiveness of utilization of added carbon source. Thereby, it is thought that a proper C:N for the biofloc system with a low salinity different from that under marine condition should be called for, in order to response the biotic or abiotic conditions at low salinity. However, information with regard to optimal C:N for biofloc system rearing *L. vannamei* under a low salinity condition has not been well documented yet.

The main aim of the present study was to estimate the optimal C:N for the biofloc system rearing *L. vannamei* at a low salinity, via investigating the effects of different C:Ns on water quality, shrimp growth during grow-out culture of *L. vannamei* at a salinity of about 5‰.

## 2. Methods and materials

### 2.1. Preparation of culture water

Water for culture experiment with a salinity of approximate 5.0‰, was prepared according to the method of Ray and Lotz (2017), with some modifications. Briefly, artificial sea salt powder (Qianglong corporation, Tianjin, China), and chemical reagents of NaHCO_3_, KCl, MgCl_2_·6H_2_O and CaCl_2_ (food grade), were added to tap water to make a salinity of ~ 5.0‰, a pH value near 8.0, and a carbonate alkalinity of about 200 mg L^-1^ CaCO_3_, with K^+^, Mg^2+^ and Ca^2+^ concentrations of about 300, 900 and 300 mg L^-1^, respectively. Before being used, sterilization with 10.0 mg L^-1^ of chlorinedioxide was conducted, followed by neutralization with 1.0 mg L^-1^ of ascorbic acid (Lara et al., 2017; Gaona et al., 2017).

### 2.2. Shrimp acclimation

*L. vannamei* shrimp juveniles raised in a culture system with a salinity of approximate 5‰, were kindly supplied from Baifuteng eco-agriculture development Co., Ltd (Changde, Hunan Province, China). Those juveniles were acclimated in the water prepared above for two weeks in the laboratory of Hunan University of Arts and Science (HUAS). During this period, juveniles were fed with a commercial formulated shrimp diet (crude protein 40.0%, crude lipid 5.0%, crude fibber 5.0%, crude ash 15.0%, moisture 12.0%, Alpha corporation, Guangdong Province, China), according to the manufacturer’s instructions. For maintaining a good water quality, feces and residual feed were siphoned out, and 20% of water was exchanged every day.

### 2.3. Experimental design and manipulations

For culture experiment, each of eighteen 500-L HDPE (high density polyethylene) tank was filled with 0.3 m^3^ of culture water prepared above, and aerated continuously with a 750-w power of air pump (HG-750-S, Sunsun Group, Zhoushan, Zhejiang, China). The tanks were randomly and equally divided into three treatments with an inputted C:N of 8:1 (CN8), 16:1 (CN16) and 24:1 (CN24), respectively. The inputted C:N was the C:N contained in the inputted materials (feed and carbon source), and determined basing on the feed carbon content (Kumar et al., 2017), and the assumption that 25% feed nitrogen would be theoretically converted as biomass and the rest (75%) lost to water body (Piedrahita, 2003).

After acclimation, shrimp juveniles (~ 0.8 g) were transferred randomly to each tank at a density of 270 shrimp m^-3^. During the 63-days culture experiment, shrimp were fed with the commercial diet same to that used during acclimation period three times a day at 8:00, 16:00 and 22:00 equivalently, basing on the estimated biomass and the feeding table according to the method by Van Wyk et al. (1999). As well, glucose (food grade, carbon content 36%, Fufeng biotechnology Co., Ltd., Hohhot, Inner Mongolia Autonomous, China) was added as the exogenous carbon source, according to the inputted C:N assigned for each treatment. No water exchange was operated for all tanks throughout the whole experiment period, except that evaporating water was complemented with dechlorinated tap water per week.

### 2.3. Water quality monitoring and measurement

Water temperature, dissolved oxygen (DO) and pH were detected *in situ* each day, using electric analyzers (YSI-ProPlus, Yellow Springs Instruments Inc., USA; pH-100 meter, LICHEN Sci-Tech, Co., Ltd., Shanghai, China). For determination of other water indices, water was sampled once a week and passed thought 0.45 μm glass fiber microfilter (Xinya purification equipment Co., Ltd, Shanghai, China). Then, total ammonia nitrogen (TAN), nitrite, nitrate, carbonate alkalinity and total suspended solid (TSS), were measured according to the standard methods (APHA, 1995). Biofloc volume (settleable solids) was determined each week with an Imhoff cone, by reading the sediment volume after a 15-min settlement of 1 L water sample (Avnimelech, 2015). The sludge volume index (SVI) was defined as the volume in milliliters occupied by 1 g of TSS after settling (Liu et al., 2014). The level of free type ammonia (NH_3_) was calculated according to the formula of Mosquera-Corral et al. (2005):

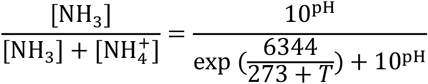

wherein, *T* was the water temperature.

### 2.4. Measurement of growth performance

At harvest, all shrimp were counted, and individually weighed with an electric balance (UW2200H, Shimadzu, Japan). And weekly increment of body weight (wiW), specific growth rate (SGR), feed conversion ratio (FCR), productivity and survival rate, were calculated according to the formulates as follows:

wiW (g week^−1^) = (fbw – ibw)/culture weeks
SGR (% d^−1^) = [(ln fbw – ln ibw)/culture days] × 100%
FCR = feed mass/(harvested biomass – stocking biomass)
Productivity (kg m^−3^) = harvested biomass/culture water volume
Survival rate (%) = surviving counts/stocking counts × 100%

Wherein, fbw and ibw represented the final and initial mean body weight of shrimp, respectively.

### 2.5. Statistical analysis

One-way ANOVA was executed with the SPSS software (version 22.0, IBM Co., NY, USA), as soon as normality distribution and variances homogeneity of data were proved with Shapiro-Wilk’s and Levene’s test, respectively. Tukey test was adopted for post hoc multiple comparisons if there was significant difference among treatments. Or else, non-parameter Kruskal-Wallis test was conducted for significant difference detecting among data of other parameters, such as pH, carbonate alkalinity and TAN. Percentage data was submitted to arcsine transformation before statistical analyses or tests. Spearman correlation coefficients between parameters were analyzed. Linear, Quadratic, Growth, Logarithmic, Cubic, Exponential, Power and Logistic models were used for curve regression and optimal C:N estimation. Differences were considered significant when *P* < 0.05.

## 3. Results

### 3.1. Water quality

The level of dissolved oxygen and water temperature were above 5.0 mg L^-1^ and 28.0 °C, respectively, in the present study. The dissolved oxygen levels in CN8 treatment was significantly higher than those in the other two treatments (*P* < 0.05, Table 1).

**Table 1.** Water parameters in the three biofloc systems with a carbon to nitrogen ratio (C:N, contained in the inputted feed and carbon source with the assumption that 75% of the feed nitrogen is excreted) of 8:1 (CN8), 16:1 (CN16) and 24:1 (CN24), respectively, during the 63-days experiment rearing *Litopenaens vannamei* at a salinity of 5.0‰

| Parameters | CN8 | CN16 | CN24 |
| --- | --- | --- | --- |
| Dissolved oxygen (mg L<sup>-1</sup>) | 5.57 ± 0.10<sup>b</sup> | 5.39 ± 0.07<sup>a</sup> | 5.28 ± 0.11<sup>a</sup> |
| Water temperature (°C) | 28.8 ± 0.3 | 28.7 ± 0.2 | 28.7 ± 0.1 |
| pH | 6.9 ± 0.1<sup>a</sup> | 7.6 ± 0.3<sup>b</sup> | 7.7 ± 0.1<sup>b</sup> |
| CAK (mg L<sup>-1</sup> CaCO<sub>3</sub>) | 104.0 ± 2.8<sup>a</sup> | 157.6 ± 21.6<sup>b</sup> | 173.0 ± 20.4<sup>b</sup> |
| TAN (mg L<sup>-1</sup>) | 7.1 ± 0.9<sup>b</sup> | 2.0 ± 0.6<sup>a</sup> | 1.3 ± 0.2<sup>a</sup> |
| NH<sub>3</sub> (mg L<sup>-1</sup>) | 0.027 ± 0.004 | 0.037 ± 0.005 | 0.029 ± 0.006 |
| Nitrite (mg L<sup>-1</sup>) | 14.0 ± 3.6<sup>b</sup> | 4.9 ± 3.1<sup>a</sup> | 1.9 ± 2.1<sup>a</sup> |
| Nitrate (mg L<sup>-1</sup>) | 77.0 ± 5.0<sup>b</sup> | 14.7 ± 5.9<sup>a</sup> | 4.6 ± 2.9<sup>a</sup> |
| Biofloc volume (mL L<sup>-1</sup>) | 4.8 ± 0.9<sup>a</sup> | 24.1 ± 3.7<sup>b</sup> | 30.5 ± 6.1<sup>b</sup> |
| TSS (mg L<sup>-1</sup>) | 327.4 ± 24.4<sup>a</sup> | 508.1 ± 32.3<sup>b</sup> | 682.1 ± 63.1<sup>b</sup> |
| SVI (mL g<sup>-1</sup>) | 11.1 ± 1.7<sup>a</sup> | 74.9 ± 5.2<sup>b</sup> | 55.1 ± 6.3<sup>b</sup> |

Note: CAK, carbonate alkalinity; SVI, sludge volume index; TAN, total ammonia nitrogen; TSS, total suspended solids. Data within a parameter with different low-case letters were significantly different ( $P < 0.05$ ).

The pH and carbonate alkalinity in CN16 and CN24 treatments were 7.6±0.3 and 157.6±21.6 mg L^-1^ CaCO_3_, and 7.7±0.1 and 173.0±20.4 mg L^-1^ CaCO_3_, respectively, which were significantly higher than those in CN8 treatment (6.9±0.1 and 104.0±2.8 mg L^-1^ CaCO_3_, *P* < 0.05, Table 1). After 28 d, the pH and carbonate alkalinity abruptly decreased in CN8 treatment, but were stable in the other two treatments throughout the whole experimental period (Fig. 1).

**Fig. 1.**
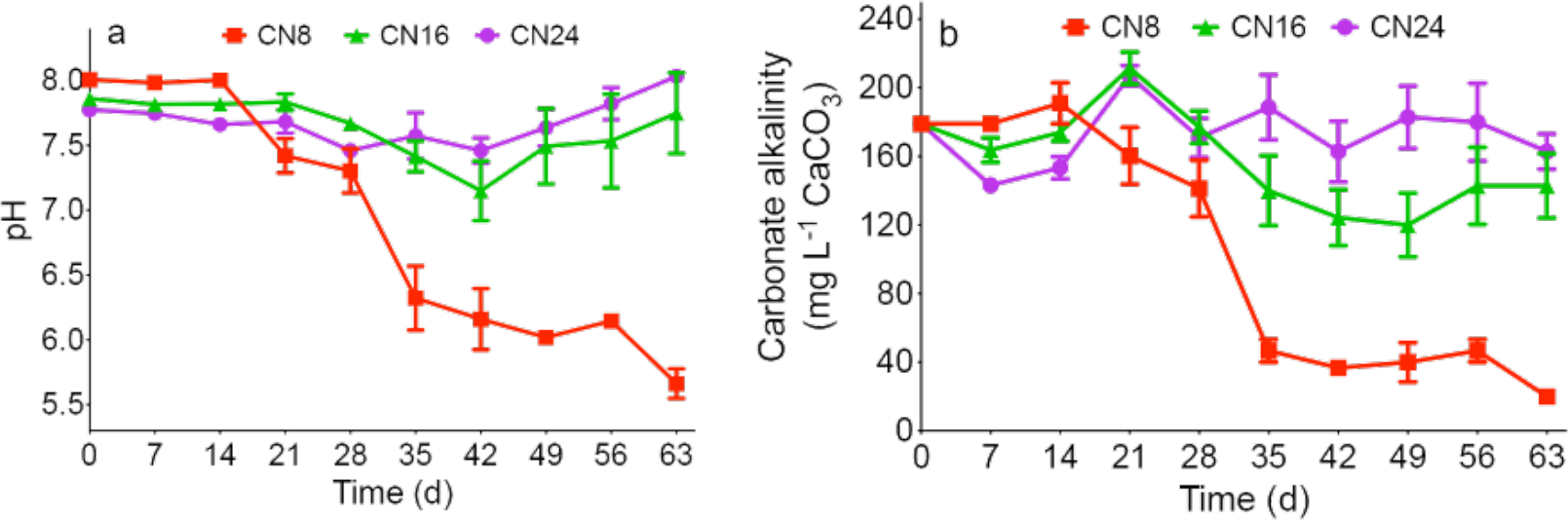
The levels of pH (a) and carbonate alkalinity (b) determined throughout the 63-days experimental period in the three biofloc treatments rearing *Litopenaeus vannamei* at a salinity of 5.0‰ with a carbon to nitrogen ratio (C:N, contained in the inputted feed and carbon source with the assumption that 75% of the feed nitrogen is excreted) of 8:1 (CN8), 16:1 (CN16) and 24:1 (CN24). Error bar indicates ± standard deviation (SD).

The TAN levels in CN16 and CN24 treatments were 2.0±0.6 and 1.3±0.2 mg L^-1^, and significantly lower than that of CN8 treatment (7.1±0.9 mg L^-1^, *P* < 0.05, Table 1). The levels of the unionized type of ammonia, NH_3_, were not significantly different among the three treatments, with a mean level below 0.037±0.005 mg L^-1^ (*P* > 0.05, Table 1). The mean nitrite level in CN8 treatment was 14.0±3.6 mg L^-1^, significantly higher than those of CN16 and CN24 treatments (4.9±3.1 and 1.9±2.1 mg L^-1^, *P* < 0.05, Table 1). Similarly, the mean nitrate level in CN8 treatment reached 77.0±5.0 mg L^-1^, far higher than those of CN16 and CN24 treatments (14.7±5.9 and 4.6±2.9 mg L^-1^, *P* < 0.05, Table 1). The TAN, NH_3_ and nitrite levels in the three treatments peaked sequentially and then decreased (Fig. 2 a-c). However, TAN in CN8 treatment accumulated again after achievement to the lowest level (Fig. 2 a). The nitrate level showed obvious, moderate and little accumulation in the CN8, CN16 and CN24 treatment, respectively (Fig. 2 d).

**Fig. 2.**
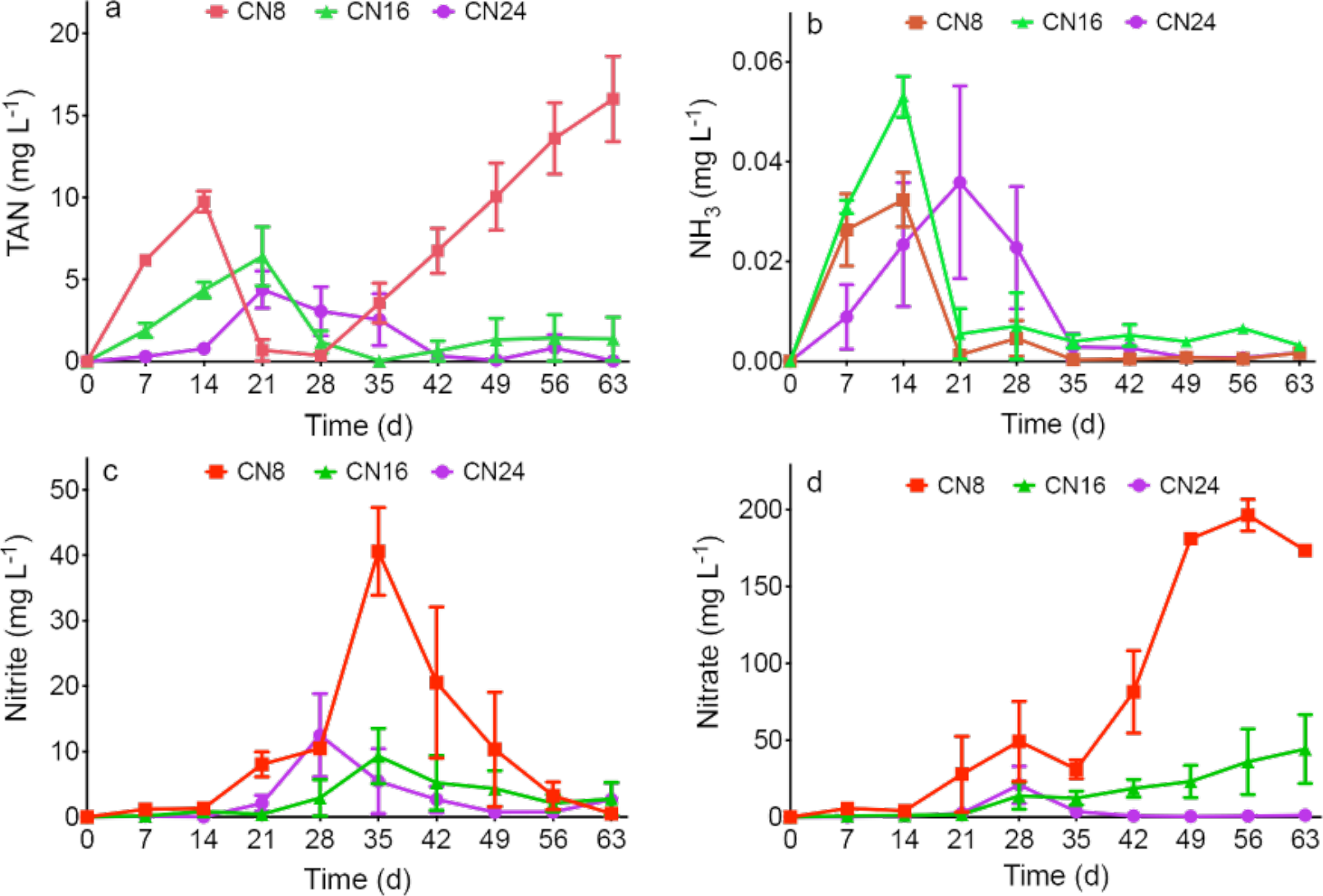
TAN (total ammonia nitrogen, a), nitrite (b) and nitrate (c) concentrations detected throughout the 63-days experimental period in the three biofloc treatments rearing *Litopenaeus vannamei* at a salinity of 5.0‰ with a carbon to nitrogen ratio (C:N, contained in the inputted feed and carbon source with the assumption that 75% of the feed nitrogen is excreted) of 8:1 (CN8), 16:1 (CN16) and 24:1 (CN24). Error bar indicates ± standard deviation (SD).

The levels of biofloc volume and TSS in CN16 and CN24 treatments were above 24 mL L^-1^ and 500 mg L^-1^, respectively, and significantly higher than those in CN8 treatment (4.8±0.9 mL L^-1^ and 327.4±24.4 mg L^-1^, *P* < 0.05, Table 1). The sludge volume index (SVI) in CN8 treatment was only 11.1±1.7 mL g^-1^, significantly deviation (SD). lower than those in CN16 and CN24 treatments (74.9±5.2 and 55.1±6.3 mL g^-1^, *P* < 0.05, Table 1). Biofloc volume and TSS accumulated throughout the experiment in the three treatments, except the biofloc volume in CN8 treatment (Fig. 3).

**Fig. 3.**
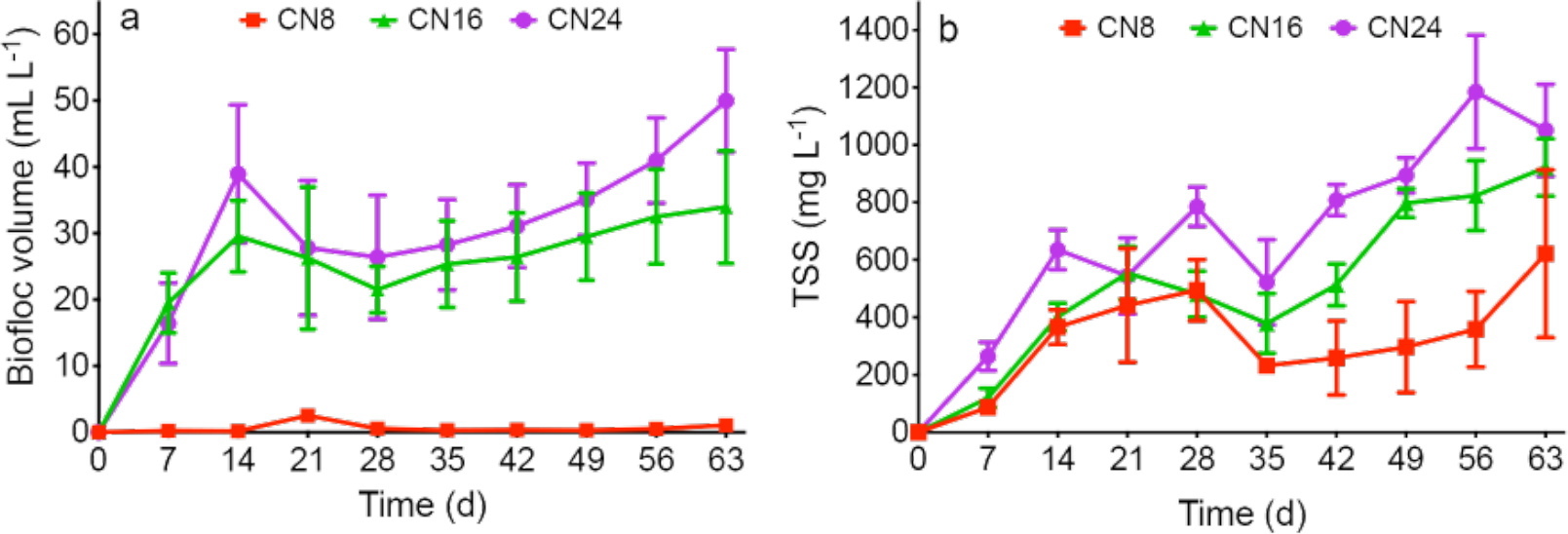
Levels of biofloc volume (a) and TSS (total suspended solids, b) measured throughout die 63-days experimental period in the three biofloc treatments rearing *Litopenaeus vannamei* at a salinity of 5.0‰ with a carbon to nitrogen ratio (C:N, contained in the inputted feed and carbon source with the assumption that 75% of the feed nitrogen is excreted) of 8:1 (CN8), 16:1 (CN16) and 24:1 (CN24). Error bar indicates ± standard deviation (SD).

The regression curves of CAK, BFV, TSS, DO, nitrite, pH, SVI, TAN and nitrate to C:N were significant (R^2^ = 0.241~ 0.846, *P* < 0.05, Table 2). Among those parameters, the first three increased (Fig. 4), but the next two decreased (Fig. 5) with increasing of C:N. The optimal C:N for the last four parameters were 20.5:1, 18.5:1, 21.0:1 and 20.8:1, respectively (Fig. 6).

**Table 2.** Statistical information of fit curves of water or growth parameters significantly regressed to the carbon to nitrogen ratio (C:N, contained in the inputted feed and carbon source with the assumption that 75% of the feed nitrogen is excreted) during the 63-days experiment rearing *Litopenaens vannamei* at a salinity of 5.0‰

| Parameter | Model type | formula | R<sup>2</sup> | P-value |  |  |
| --- | --- | --- | --- | --- | --- | --- |
|  |  |  |  | model | Coefficient |  |
| TAN | Quadratic | $y=17.5-1.6x+0.04x^2$ | 0.784 | < 0.001 | Constant | < 0.001 |
|  |  |  |  |  | x | < 0.001 |
|  |  |  |  |  | x<sup>2</sup> | 0.001 |
| CAK | Power | $y=47.9x^{0.41}$ | 0.604 | < 0.001 | Constant | 0.001 |
|  |  |  |  |  | x | < 0.001 |
| Nitrate | Quadratic | $y=201.2-19.2x+0.46x^2$ | 0.846 | < 0.001 | Constant | < 0.001 |
|  |  |  |  |  | x | 0.001 |
|  |  |  |  |  | x<sup>2</sup> | 0.003 |
| pH | Quadratic | $y=5.4+0.2x-0.006x^2$ | 0.645 | 0.001 | Constant | < 0.001 |
|  |  |  |  |  | x | 0.002 |
|  |  |  |  |  | x<sup>2</sup> | 0.006 |
| BFV | Growth | $y=e^{(1.5+0.08x)}$ | 0.377 | 0.009 | Constant | 0.008 |
|  |  |  |  |  | x | 0.009 |
| SVI | Quadratic | $y=-140.1+24.1x-0.65x^2$ | 0.754 | < 0.001 | Constant | < 0.001 |
|  |  |  |  |  | x | < 0.001 |
|  |  |  |  |  | x<sup>2</sup> | < 0.001 |
| DO | Logarithmic | $y=6.1-0.26\log(x)$ | 0.620 | < 0.001 | Constant | < 0.001 |
|  |  |  |  |  | x | < 0.001 |
| Nitrite | Growth | $y=e^{(3.6-0.14x)}$ | 0.241 | 0.045 | Constant | 0.009 |
|  |  |  |  |  | x | 0.045 |
| TSS | Power | $y=93.3x^{0.61}$ | 0.622 | < 0.001 | Constant | 0.012 |
|  |  |  |  |  | x | < 0.001 |
| SR | Quadratic | $y=57.9+4.3x-0.12x^2$ | 0.383 | 0.034 | Constant | < 0.001 |
|  |  |  |  |  | x | 0.016 |
|  |  |  |  |  | x<sup>2</sup> | 0.022 |

Note: BFV, biofloc volume; CAK, carbonate alkalinity; DO, dissolved oxygen; SR, survival rate; SVI, sludge volume index; TAN, total ammonia nitrogen; TSS, total suspended solids. Data within a parameter with different low-case letters were significantly different ( $P < 0.05$ ).

**Fig. 4.**
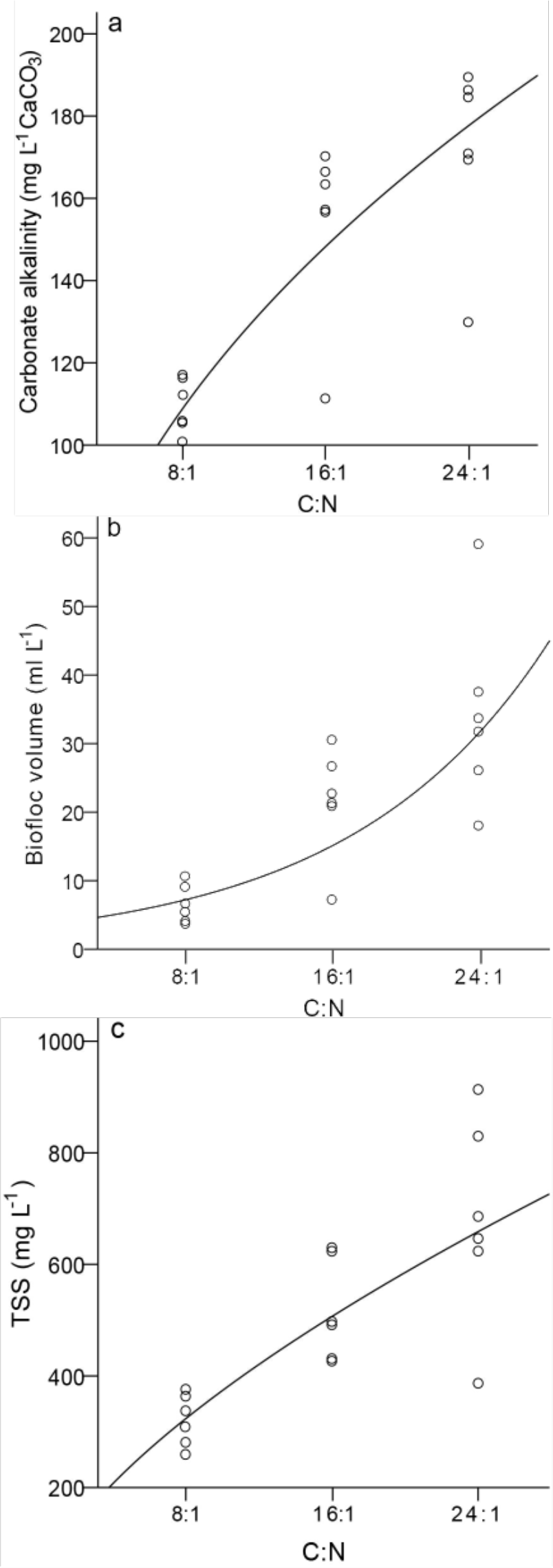
Scatterplots and fit curves of carbonate alkalinity (a), biofloc volume (b) and TSS (c) to the carbon to nitrogen ratio (C:N, contained in the inputted feed and carbon source with the assumption that 75% of the feed nitrogen is excreted) during the 63-days experiment rearing *Litopenaeus vannamei* at a salinity of 5.0‰. TSS, total suspended solids.

**Fig. 5.**
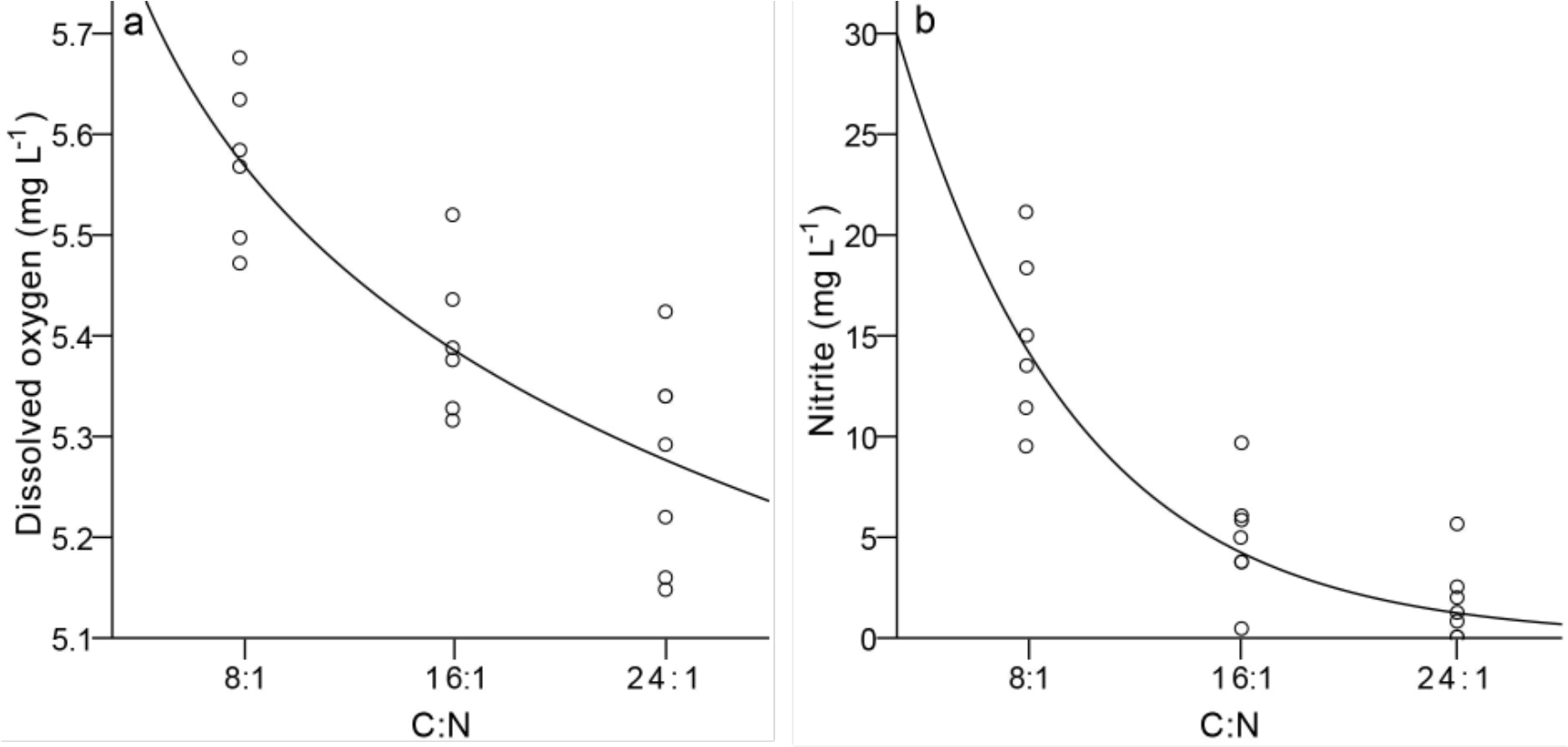
Scatterplots and fit curves of dissolved oxygen (a) and nitrite (b) to the carbon to nitrogen ratio (C:N, contained in the inputted feed and carbon source with the assumption that 75% of the feed nitrogen is excreted) during the 63-days experiment rearing *Litopenaeus vannamei* at a salinity of 5.0‰.

**Fig. 6.**
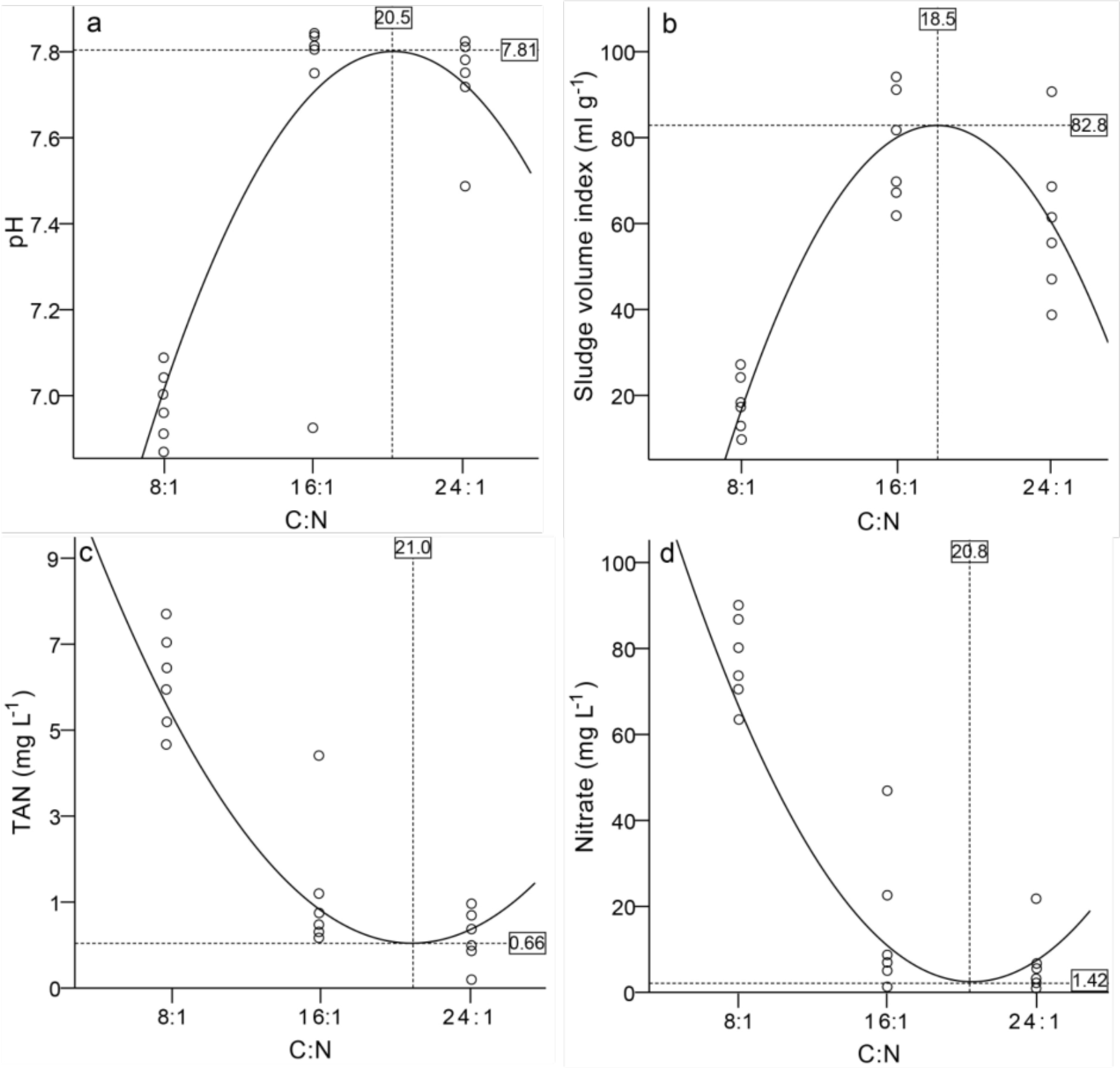
Scatterplots and fit curves of pH (a), sludge volume index (b), TAN (c) and nitrate (d) to the carbon to nitrogen ratio (C:N, contained in the inputted feed and carbon source with the assumption that 75% of the feed nitrogen is excreted) during the 63-days experiment rearing *Litopenaeus vannamei* at a salinity of 5.0‰. TAN. total ammonia nitrogen.

The correlations between the biofloc-associating parameters (such as BFV, TSS and SVI) and the other water parameters were estimated in the present study. BFV and TSS negatively correlated with DO and the three inorganic nitrogen compounds (r = −0.542 ~ −0.770, *P* < 0.05), but positively with CAK (r = 0.676 and 0.789, *P* < 0.05, Table 3). BFV also positively correlated with pH (r = 0.551, *P* < 0.05, Table 3). In regard to SVI, it only significantly correlated with pH and TAN (r = 0.495 and −0.525, *P* < 0.05, Table 3).

**Table 3.** Spearman correlation coefficients between parameters related to biofloc and the other water parameters during the 63-days experiment rearing *Litopenaeus vannamei* at a salinity of 5.0‰

|  | DO | Temp | pH | CAK | TAN | NH<sub>3</sub> | Nitrite | Nitrate |
| --- | --- | --- | --- | --- | --- | --- | --- | --- |
| BFV | -0.542* | -0.033 | 0.551* | 0.676** | -0.706** | 0.132 | -0.770** | -0.740** |
| TSS | -0.574* | 0.209 | 0.429 | 0.789** | -0.554* | 0.201 | -0.767** | -0.755** |
| SVI | -0.299 | -0.344 | 0.495* | 0.167 | -0.525* | -0.005 | -0.243 | -0.314 |

Note: BFV, biofloc volume; CAK, carbonate alkalinity; DO, dissolved oxygen; SVI, sludge volume index; TAN, total ammonia nitrogen; Temp, water temperature; TSS, total suspended solids. BFV, TSS and SVI were the parameters related to biofloc. \*, $P < 0.05$ ; \*\*, $P < 0.01$ .

### 3.2. Growth performance of shrimp

The survival rates in CN16 and CN24 treatments were 96.8±2.0% and 93.7±4.2%, respectively, which were significantly higher than that of CN8 treatment (81.5±6.4%, *P* < 0.05, Table 4). There was no significant difference on the other determined zootechnical parameters of shrimp among three treatments (*P* > 0.05, Table 4). The final mean body weight of shrimp in the present study was 11.27±0.83 ~ 12.46±1.02 g, with a weekly increment of body weight (wiW) of 1.14±0.25 ~ 1.21±0.08 g week^-1^ (Table 4). The productivity were 2.73±0.12, 3.06±0.08 and 2.82±0.18 kg m^-3^ in the CN8, CN16 and CN24 treatment, respectively (*P* > 0.05, Table 4).

**Table 4.** Growth performance of *Litopenaeus vannamei* cultured in the three biofloc systems with a carbon to nitrogen ratio (C:N, contained in the inputted feed and carbon source with the assumption that 75% of the feed nitrogen is excreted) of 8:1 (CN8), 16:1 (CN16) and 24:1 (CN24), respectively, during the 63-days experiment rearing *Litopenaeus vannamei* at a salinity of 5.0‰

| Zootechnical parameters | CN8 | CN16 | CN24 |
| --- | --- | --- | --- |
| Initial body weight (g) | $0.79 \pm 0.04$ | $0.83 \pm 0.08$ | $0.82 \pm 0.07$ |
| Final body weight (g) | $12.46 \pm 1.02$ | $11.71 \pm 0.29$ | $11.27 \pm 0.83$ |
| wiW ($\text{g week}^{-1}$) | $1.19 \pm 0.10$ | $1.21 \pm 0.08$ | $1.14 \pm 0.25$ |
| SGR ($\% \text{ d}^{-1}$) | $4.24 \pm 0.13$ | $4.20 \pm 0.16$ | $4.10 \pm 0.25$ |
| FCR | $1.24 \pm 0.10$ | $1.21 \pm 0.08$ | $1.34 \pm 0.36$ |
| Survival rate (%) | $81.5 \pm 6.4^a$ | $96.8 \pm 2.0^b$ | $93.7 \pm 4.2^b$ |
| Productivity ($\text{kg m}^{-3}$) | $2.73 \pm 0.12$ | $3.06 \pm 0.08$ | $2.82 \pm 0.18$ |

Note: FCR, feed conversion ratio; SGR, specific growth rate; wiW, weekly increment of body weight. Data within a parameter with different low-case letters were significantly different ( $P < 0.05$ ).

Among the growth performance parameters, only survival rate was significantly regressed to C:N (R^2^ = 0.383, *P* = 0.034, Table 2), with an optimal C:N of 18.6:1 (Fig. 7).

**Fig. 7.**
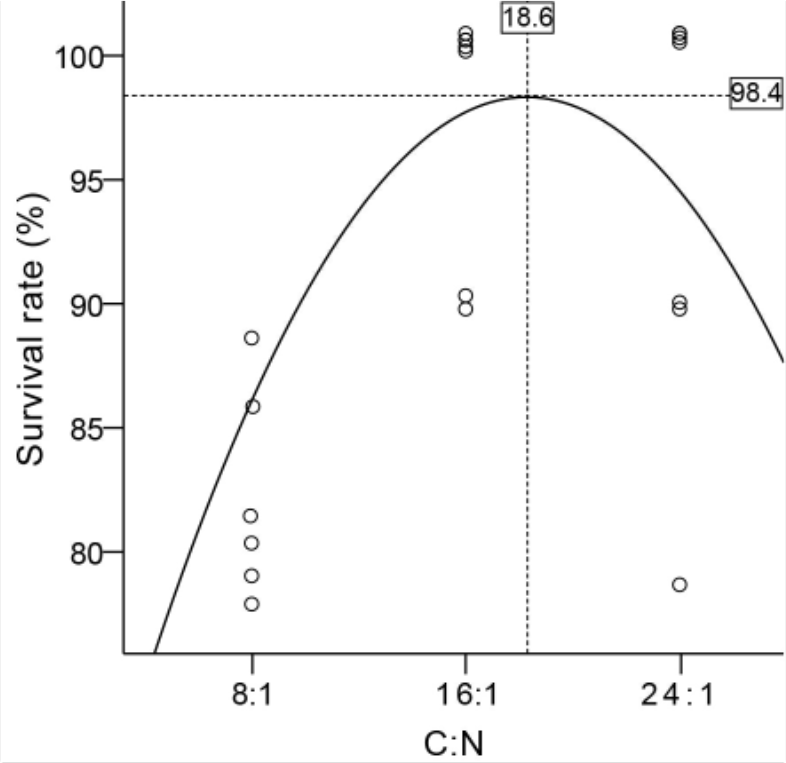
Scatterplots and fit curve of survival rate of shrimp to the carbon to nitrogen ratio (C:N, contained in the inputted feed and carbon source with the assumption that 75% of the feed nitrogen is excreted) during the 63-days experiment rearing *Litopenaeus vannamei* at a salinity of 5.0‰.

The results of correlation analysis showed that there was no significant correlation between dissolved oxygen, water temperature, TAN, the free-type ammonia (NH_3_) with zootechnical parameters of shrimp (*P* > 0.05, Table 5). However, nitrite significantly negatively correlated with weekly increment of body weight (wiW, r = −0.596, *P* < 0.05), survival rate (SR, r = −0.504, *P* < 0.05) and productivity (Prod, r = −0.597, *P* < 0.05), and positively with FCR (r = 0.597, *P* < 0.05, Table 5). In addition, significant correlation coefficients between survival rate and pH (r = 0.535), carbonate alkalinity (CAK, r = 0.598), nitrate (r = −0.567), biofloc volume (BFV, r = 0.598), TSS (r = 0.630) and SVI (r = 0.535) were found (*P* < 0.05, Table 5).

**Table 5.** Correlation coefficients (Spearman) between water parameters and shrimp zootechnical parameters during the 63-days experiment rearing *Litopenaeus vannamei* at a salinity of 5.0‰

|  | fbW | wiW | SGR | FCR | SR | Prod |
| --- | --- | --- | --- | --- | --- | --- |
| DO | 0.383 | 0.221 | 0.221 | -0.208 | -0.315 | 0.208 |
| Temp | -0.052 | 0.042 | 0.228 | -0.043 | 0.047 | 0.043 |
| pH | 0.022 | 0.260 | 0.098 | -0.262 | 0.535* | 0.262 |
| CAK | 0.148 | 0.480 | 0.088 | -0.472 | 0.598* | 0.472 |
| TAN | 0.080 | -0.120 | 0.118 | 0.121 | -0.378 | -0.121 |
| NH<sub>3</sub> | 0.156 | 0.319 | 0.287 | -0.320 | -0.126 | 0.320 |
| Nitrite | -0.330 | -0.596* | -0.331 | 0.597* | -0.504* | -0.597* |
| Nitrate | -0.058 | -0.360 | 0.022 | 0.362 | -0.567* | -0.362 |
| BFV | 0.036 | 0.346 | 0.368 | -0.347 | 0.598* | 0.347 |
| TSS | -0.178 | 0.150 | 0.015 | -0.146 | 0.630** | 0.146 |
| SVI | -0.224 | -0.012 | -0.096 | 0.009 | 0.535* | -0.009 |

Note: BFV, biofloc volume; CAK, carbonate alkalinity; DO, dissolved oxygen; fbW, final body weight; FCR, feed conversion rate; Prod, productivity; SGR, specific growth rate; SR, survival rate; SVI, sludge volume index; TAN, total ammonia nitrogen; Temp, water temperature; TSS, total suspended solids; wiW, weekly increment of body weight. \*, $P < 0.05$ ; \*\*, $P < 0.01$ .

## 4. Discussion

### 4.1. Influence of C:N on water quality

The water parameters were all in the acceptable range for culture of shrimp in the present study (Furtado et al., 2014; Hargreaves, 2013; Schveitzer et al., 2013; Van Wyk et al., 1999; Xu et al., 2012; Zhang et al., 2017), except that in the CN8 treatment, pH and biofloc volume were lower than the recommend levels (Samocha et al., 2007; Van Wyk et al., 1999), and that TAN and nitrite were higher than their safety levels (Lin and Chen, 2001; Lin and Chen, 2003). Previous studies showed that water parameters vibrated in biofloc systems along with increasing of C:N (Chakrapani et al., 2021; Panigrahi et al., 2019; Panigrahi et al., 2018; Xu et al., 2016; Xu et al., 2018). Xu et al. (2016) assumed that with the increase in C:N (from 9:1 to 18:1), there was a shift in the dominant microbial community from the original mix of photoautotrophic microalgae and chemoautotrophic bacteria to predominantly chemoautotrophic bacteria and further to heterotrophic bacteria. In other words, the lower or higher carbon input would contribute to a higher production of autotrophic or heterotrophic bacteria, respectively. Consistently, predominant chemoautotrophic bacteria in the CN8 treatment, co-occurrence of chemoautotrophic and heterotrophic bacteria in CN16 treatment, and predominant heterotrophic bacteria in CN24 treatment were inferred in the present study, being corroborated by the large, moderate and few accumulation of nitrate in those treatments, respectively. Nitrate is the last product of nitrification process implemented by chemoautotrophic nitrifiers (Ebeling et al., 2006), such as ammonia-oxidizing bacteria (AOB) and nitrite-oxidizing bacteria (NOB). Thereby, large accumulation of nitrate means establishment of full nitrification, and the predominance of AOB and NOB (Luo et al., 2020).

Compared to autotrophic bacteria, heterotrophic bacteria have different characteristics, such as approximate 10 times higher growth rate, 40 times greater microbial biomass yield, and faster conversion rate of dissolved nitrogen into bioflocs with less consumption of carbonate alkalinity (HCO_3_^-^), leading to different impacts on water parameters from those of autotrophic bacteria (Avnimelech, 1999; Ebeling et al., 2006; Hargreaves, 2006; Huang, 2020). Briefly, with increasing of C:N, the growth of heterotrophic bacteria are prompted, resulting in that the levels of TAN, nitrite and nitrate decrease, but the biofloc volume and TSS levels increase (Chakrapani et al., 2021; Panigrahi et al., 2019; Panigrahi et al., 2018), in agreement with the results of the present study. Furthermore, the consumption of carbonate alkalinity would be less under higher C:N condition, promising the result of high level of carbonate alkalinity in CN16 and CN24 treatments of the present study, as well as the pH, because in aquaculture waterbody, carbonate alkalinity is very important to the buffer ability of water column to calibrate and stabilize the pH (Furtado et al., 2015a; Pierri et al., 2015). Additionally, the dissolved oxygen level will also decrease with the increasing of C:N in the biofloc system, being contributed to that respiration of thrived-growing heterotrophic bacteria and oxidation of massive-accumulated organic matter (such as biofloc, residual feed and feces) will consume a large amount of dissolved oxygen (Azhar et al., 2020). In the present study, biofloc volume and TSS significantly elevated with increasing of C:N, both of which showed negative correlations with the dissolved oxygen level.

It is worthy of note that TAN in CN8 treatment accumulated again after achievement to the lowest level at 28 d, indicating that it might be imbalanced among activities of ammonia producing, ammonia oxidizing and nitrite oxidizing in this treatment. However, the details should be studied next.

Partial water parameters in the present study were found to be significantly regressed with C:N. However, the effects of C:N on water parameters weakened gradually, making us induce an optimal C:N range of 18.5:1-21.0:1. This is the first time to quantitatively investigate the relationship between C:N and water quality for the biofloc system under a low salinity condition, to the best of our knowledge. However, it was also found in the present study that a C:N of 16:1 was enough to maintain a good water quality and obtain a good growth performance. Previously, a C:N range of 10:1-20:1 was considered to be optimal to management of water quality for the marine biofloc systems (Avnimelech, 1999; Tong et al., 2020).

### 4.2. Influence of C:N on shrimp growth performance

In some previous studies, the growth performance of shrimp is elevated with increasing of C:N (Chakrapani et al., 2021; Panigrahi et al., 2019; Panigrahi et al., 2018). Contradictorily, in other studies, the shrimp growth performance was found to be better in biofloc systems with low C:N (Xu et al., 2016; Xu et al., 2018). Different experimental conditions, such as the indoor condition and the long 120-days experiment duration for the former studies, and the outdoor condition and the short 42-days experiment duration for the latter studies, might contribute the discrepancy. Interestingly, under the indoor condition and the moderate 63-days experiment period in the present study, higher final body weight and specific growth rate (SGR) of shrimp were observed in the biofloc system with a low C:N (CN8 treatment). The lower stocking density due to lower survival rate in the CN8 treatment of the present study might contribute this increment (Esparza-Leal et al., 2020). Whereas, the survival rate and productivity in the CN16 and CN24 treatments were found to be higher than those in the CN8 treatment. Microbial community activities in BFT could result in fluctuations of water parameters, such as pH, dissolved oxygen and alkalinity, subsequently affecting the growth and survival rate of culture organisms (Furtado et al., 2011; Khoa et al., 2020; Martins et al., 2003; Martins et al., 2017). In the current study, the authors also speculated that first, elevation of C:N directly prompts growth of heterotrophic bacteria, leading to formation and accumulation of biofloc and TSS. Second, bioactivities of heterotrophic bacteria adhering to accumulated biofloc manages the other water parameters, such as inorganic nitrogen compounds, pH, carbohydrate alkalinity and dissolved oxygen. And next, those water parameters affect the zootechnical performance of shrimp. Correspondingly, in the present study, the two parameters related to biofloc, biofloc volume (BFV) and TSS, were directly found to be positively regressed to C:N. Next although without significant correlation with the zootechnical parameters (except the survival rate), BFV and TSS, the first two parameters directly affected by C:N, were found to significantly correlate with some other water parameters, such as pH, carbonate alkalinity, nitrite and nitrate. And subsequently, among those water parameters, nitrite showed significant correlation coefficients with the zootechnical parameters, such as weekly increment of body weight (wiW), feed conversion ratio (FCR), survival rate (SR) and productivity (Prod). As well, pH, carbonate alkalinity and nitrate were all significantly correlate with the survival rate. Those findings indicated that C:N indirectly affected the growth performance of shrimp in the present study.

The zootechnical parameters of shrimp in CN16 treatment were better than those in CN24 in the present study (*P* > 0.05). This might be contributed to the dominant mixture of autotrophic and heterotrophic bacteria in CN16 treatment, but dominant heterotrophic bacteria in CN24 treatment, being inferred by the moderate and low levels of nitrate in both treatments, respectively. While the dominated mixture bacteria in biofloc systems are considered to be more beneficial for the growth of shrimps than the dominated heterotrophic bacteria (Khoa et al., 2020; Xu et al., 2016).

### 4.3. Influence of water quality on shrimp growth performance

The nitrite in the present study showed negative correlations with weekly increment of body weight (wiW), survival rate (SR) and productivity (Prod). Nitrite binds to hemocyanin, causes oxidation of copper atoms (Cu^+^ to Cu^2+^) such that the functional hemocyanin molecule is converted into meta-hemocyanin which cannot reversibly bind molecular oxygen (Chen and Chin, 1988; Ferreira et al., 2020), preventing the transport of oxygen to tissues and reducing the amount of oxygen available for metabolism (Tahon et al., 1988). This process can lead to hypoxia and, consequently, mortality of organisms cultured (Morais et al., 2020). Though nitrate is less toxicity to animals (Furtado et al., 2015b), it also showed negative correlations with survival rate in this study. This relationship might be contributed to its close association with nitrite. Nitrate is the end product of nitrification process produced from nitrite (Ebeling et al., 2006), indicating that its level is related to that of nitrite, and that high level of nitrate means that high level of nitrite could have been inferred. And thus, the nitrate level would be subsequently related to the levels of the parameters correlated with nitrite. Whereas, in the present study, TAN did not show significant effect on growth parameters of juveniles. TAN, the total ammonia nitrogen, contains two types, ionic type (NH_4_^+^) and unionized type (NH_3_); and the latter is the derivate of toxicity of TAN (Green et al., 2020). NH_3_ with a level of < 0.03 mg L^-1^ is considered to be safe for *L. vannamei* at the low salinity condition; whereas, chronic effects or lethality will appear if the concentration is higher than 0.1 mg L^-1^ (Van Wyk et al., 1999). In the present study, the mean level of NH_3_ was lower than 0.037±0.005 mg L^-1^, indicating a low toxicity level for shrimp.

The other water parameters did not show significant correlation with the zootechnical parameters in the present study, with exception of pH, carbonate alkalinity, biofloc volume, TSS and sludge volume index (SVI). Those parameters showed significant correlations with the survival rate of shrimp, indicating that maintenance of the suitable levels for those parameters could supply a good environment with less stress for cultured animals and reduce the mortality. De Schryver et al. (2008) thought that under a high level of SVI, biofloc has the capacity to remain suspended in the water column, thus could be easily harvested by culture species to improve the growth performance. Although the levels of SVI in CN16 and CN24 treatments in the present study (55.1-74.9 mL g^-1^) reached to the optimal level for culture of *L. vannamei* (Liu et al., 2014), no improvement for the shrimp growth performance was observed when compared to that of the CN8 treatment with a low SVI (11.1±1.7 mL g^-1^). Nevertheless, the higher level of SVI might be helpful for management of other water parameters, due to supplying more suspended biofloc as substrates for colonization of bacteria. For example, significant correlations between SVI with pH and TAN were observed in the present study.

## 5. Conclusion

Under the condition of the present study, the water parameters in the treatments with a C:N higher than 16:1 (CN16 and CN24) were all in the acceptable range for culture of shrimp. However, in the treatment with a C:N of 8:1 (CN8), the pH and biofloc volume were lower than the recommend levels, and the TAN and nitrite levels were higher than their safety levels. The survival rate of shrimp in treatments of CN16 and CN24 were higher than those in CN8. The results indicated that an inputted C:N higher than 16:1 was suitable for the biofloc system rearing *L. vannamei* with a low salinity of about 5‰, with an optimal inferred C:N range of 18.5-21.0:1 for water quality and growth performance.

## Acknowledgements

This work is financially supported by the scientific research program of the Education Department of Hunan Province, China (18B394). The authors also thank Bifuteng eco-agriculture development Co., Ltd. (Changde, Hunan Province, China) for the kindly supplying of shrimp.

